# F-53B damages cell viability leading to impaired root development in Arabidopsis

**DOI:** 10.1101/2025.05.13.653422

**Authors:** Jiayi Zhao, Wenjing Pan, Yuting Chen, Xueying Cui, Huiqi Fu, Yufeng Luo, Zitong Chen, Nan Huang, Yue Hu, Yonghua Qin, Guanghui Yu, Ziming Ren, Wenyi Wang, Xiaoning Lei, Bing Liu

## Abstract

The emerging contaminant chlorinated polyfluoroalkyl ether sulfonate (Cl-PFAES/F-53B) has been detected in many plant species, however, the impact of F-53B on the development in plants and the underpinning mechanisms remain largely unelucidated. In this study, we demonstrate by molecular biology, cytogenetics, and fluorescence microscopy mythologies that F-53B impairs cell viability leading to disrupted root development in Arabidopsis (*Arabidopsis thaliana*). We show that F-53B suppresses root length and the inhibitory effect is pronunanced at increasing concentrations. F-53B disrupts cell wall and microtubule-mediated cell plate formation and configuration in meristem cells, revealing its toxicity to cell division and cytoskeleton organization. F-53B induces cell death predominantly at the meristematic zone, however, gene exression and genetic studies indicate that F-53B does not trigger DNA double-strand breaks, implying a divergent effect on DNA stability across species. Moreover, we show that F-53B damages nuclei stability and viability, and induces Autophagy-Related 8 protein foci formation implying that it triggers autophagy-mediated cell responses. Taken together, this study provides cellular and molecular insights into the toxicity of F-53B to plant cells, which highlights its damage to agriculture and ecology security that deserve imperative environmental management actions.

**Environmental Implication:** F-53B has been detected in many plant species, which raises increasing concerns on its toxicity to plants and thus agriculture and ecology safety. However, the toxicity of F-53B to development in plants and the underpinning mechanisms remain largely unelucidated. Here, we demonstrate that F-53B damages nuclei stability and viability leading to cell death and impaired root development in the model plant species *Arabidopsis thaliana*. Our study provides cytogenetic and molecular insights into the F-53B toxicity to plant cells, which highlights imperative environmental management actions and can be referenced for developing F-53B control and management policies.

**GRAPHICAL ABSTRACT:** 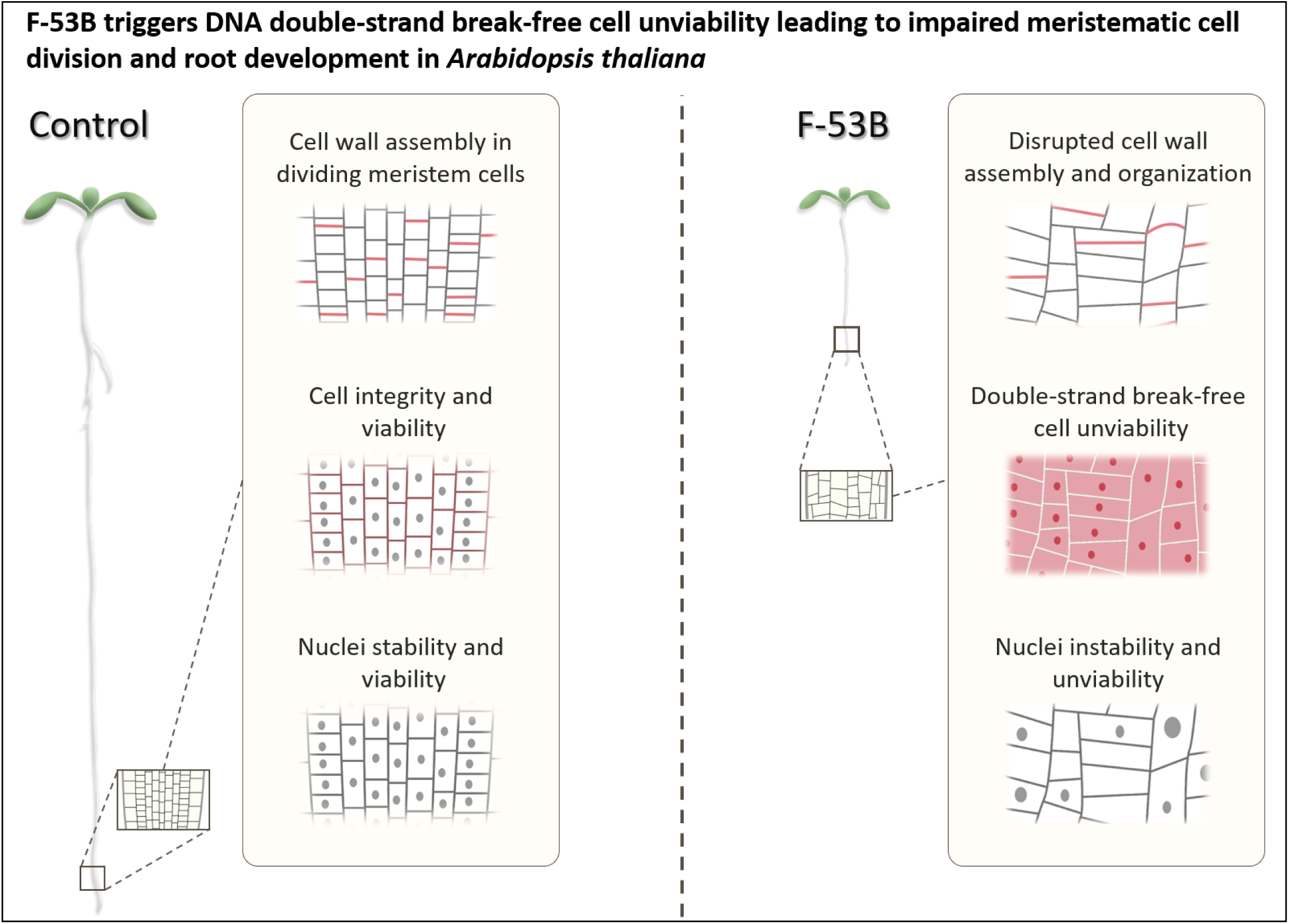

**One-sentence summary:** F-53B triggers DNA double-strand break-free cell unviability leading to impaired meristematic cell division and root development in plants.

**HIGHLIGHTS:**

- F-53B impairs root development in Arabidopsis.
- F-53B prohibits mitotic cell division at root tips.
- F-53B damages cell viability predominantly at the meristematic zone in seedlings.
- F-53B does not induce DNA double-strand breaks in Arabidopsis seedlings.
- F-53B damages nuclei viability and stability in seedlings.

## 1. Introduction

Per- and polyfluoroalkyl substances (PFAS), the so-called ‘Forever Chemicals’, are anthropogenic chemicals with strong carbon-fluorine bonds, which were manufactured in the 1950s and thereafter have been widely used in various industrial and consumer products due to their chemical and thermal stability as well as hydrophobic and hydrophilic properties (Gaines, 2023; Glüge et al., 2020). PFAS are released into environment during production, daily use, and disposal, leading to prevalent dispersion in the main environmental matrices (Brusseau et al., 2020; De Silva et al., 2021; Nakayama et al., 2019). Due to their persistence, long distance transport potential, bioaccumulation, eco-and bio-toxicity, PFAS have raised global concerns and regulatory actions especially in developed countries over the past decades (Cousins et al., 2020; Kwiatkowski et al., 2020; Ng et al., 2021; Panieri et al., 2022).

Perfluorooctane sulfonic acid (PFOS), the most common PFAS, has been detected in various environmental compartments, raising serious concerns for global crisis of PFOS contamination (Wee and Aris, 2023a, b). PFOS in irrigation water, soil and/or biosolids are absorbable by plant species including mung bean (Pan et al., 2021), maize (Ebinezer et al., 2022; Just et al., 2022), lettuce (Li et al., 2021; Yu et al., 2021), wheat (Liu et al., 2022; Ofoegbu et al., 2022; Qu et al., 2010), vegetable (Zhou et al., 2021) and Arabidopsis (Groffen et al., 2023; Kim et al., 2024; O’Hara and Longstaffe, 2022; Zhang et al., 2022). Many studies have reported that the growth and development of plants are subject to PFOS exposure. For example, PFOS affects root development, lipid peroxidation, protein carbonylation and DNA stability in multiple plant species, highlighting a threat to agricultural and ecological safety (Adu et al., 2023; Chen et al., 2020; Chi et al., 2024; Ghisi et al., 2019; He et al., 2023; Li et al., 2022; Pietrini et al., 2024; Song et al., 2024; Wang et al., 2020).

With the regulation and restriction of PFOS, emerging substitutes have been introduced into markets, in which chlorinated polyfluoroalkyl ether sulfonate (Cl-PFESA, trade name F-53B, primarily comprising 6:2 Cl-PFESA and 8:2 Cl-PFESA) has been used as the main PFOS alternative in the electroplating industry for over forty years (Wang et al., 2013). Despite F-53B is currently only used in China, it has been ubiquitously detected worldwide thus reflecting a global contamination status of F-53B (Ti et al., 2018). F-53B reveals a higher concentration than PFOS in soil, warning a serious land contamination by F-53B (Li et al., 2020). Compared with PFOS, F-53B exhibits a higher accumulation ability in plant roots, probably due to its long length of carbon chain and a sulfonic functional group (Lin et al., 2020; Xu et al., 2022; Zhang et al., 2021), greater affinity to carrier proteins and stronger phytotoxicity. Studies in mung bean and wheat have revealed that F-53B is more toxic than PFOS to plant development, possibly by triggering higher levels of hydroxyl free radicals (Li et al., 2023b; Lin et al., 2020; Pan et al., 2021). Moreover, F-53B induces a different but greater rhizosphere defense response in plants than PFOS, revealing that they have distinct impacts on plant development (Lu et al., 2023). Despite the increasing awareness of biotoxicity and thus the threats of F-53B to agriculture and ecology, the mechanism underpinning F-53B-induced development inhibition and DNA instability in plants, especially at the cellular and molecular levels, remains largely unclear.

In this study, we evaluated the inhibitory effect of F-53B on seedling development in the model plant species Arabidopsis (*Arabidopsis thaliana*) and dissected the underpinning mechanisms. We found that F-53B prohibits root development in plants by damaging nuclei stability and/or function, which leads to cell unviability and associated impaired cell division at the meristematic region at root tips. This study provides cytogenetic and molecular evidence unvealing the cell toxicity of F-53B to plants and highlights its threat to agricultural and ecological safety.

## 2. Material and Methods

### 2.1. Plant materials and growth conditions

*Arabidopsis thaliana* (L.) accession Columbia-0 (Col-0) (for simplification ‘Col’ is used throughout the manuscript) was used as the wild-type in this study. The *rad51* (GABI_134A01), *atm-2* (SALK_006953) and *mre11-3* (SALK_054418) (Puizina et al., 2004) mutants, the *pMRE11::MRE11-eGFP*, *pMRE11::GUS-eGFP*, *pRAD51::RAD51-GFP (Da Ines et al., 2013)*, *pMAP65-3::GFP-MAP65-3* (Sofroni et al., 2020), *pCENH3::RFP-CENH3* (Komaki et al., 2020), *pRPS5A::TagRFP-TUA5* (Sofroni et al., 2020), *p35S::ATG8a-GFP* (Zhou et al., 2023), *pUBQ::mCherry-ATG8e* (Zhuang et al., 2017), *pCYCB3;1::CYCB3;1-GFP* (Sofroni et al., 2020), *pCDKA;1::CDKA1-YFP* (Nowack et al., 2006) and *pPIN3::PIN3-GFP* (Wang et al., 2021) reporters were used in this study. Primers for genotyping are listed in Supplementary material Table S1. Seeds were germinated and cultivated in half-strength Murashige & Skoog (1/2 MS) medium with or without F-53B in a growth chamber with a 16 h day/8 h night, 20°C, and 50% humidity condition following vernalization in a dark and 4°C condition for 3 days. To calculate the length of main roots and lateral roots in Col, *rad51* and *atm* (Fig. 1; Fig. 7), seedlings exposed to F-53B for 8 and 14 days, respectively, were measured. For histochemical staining and live-imaging assays, seedlings exposed to F-53B for 5 days were used for experiments.

**Fig. 1.**
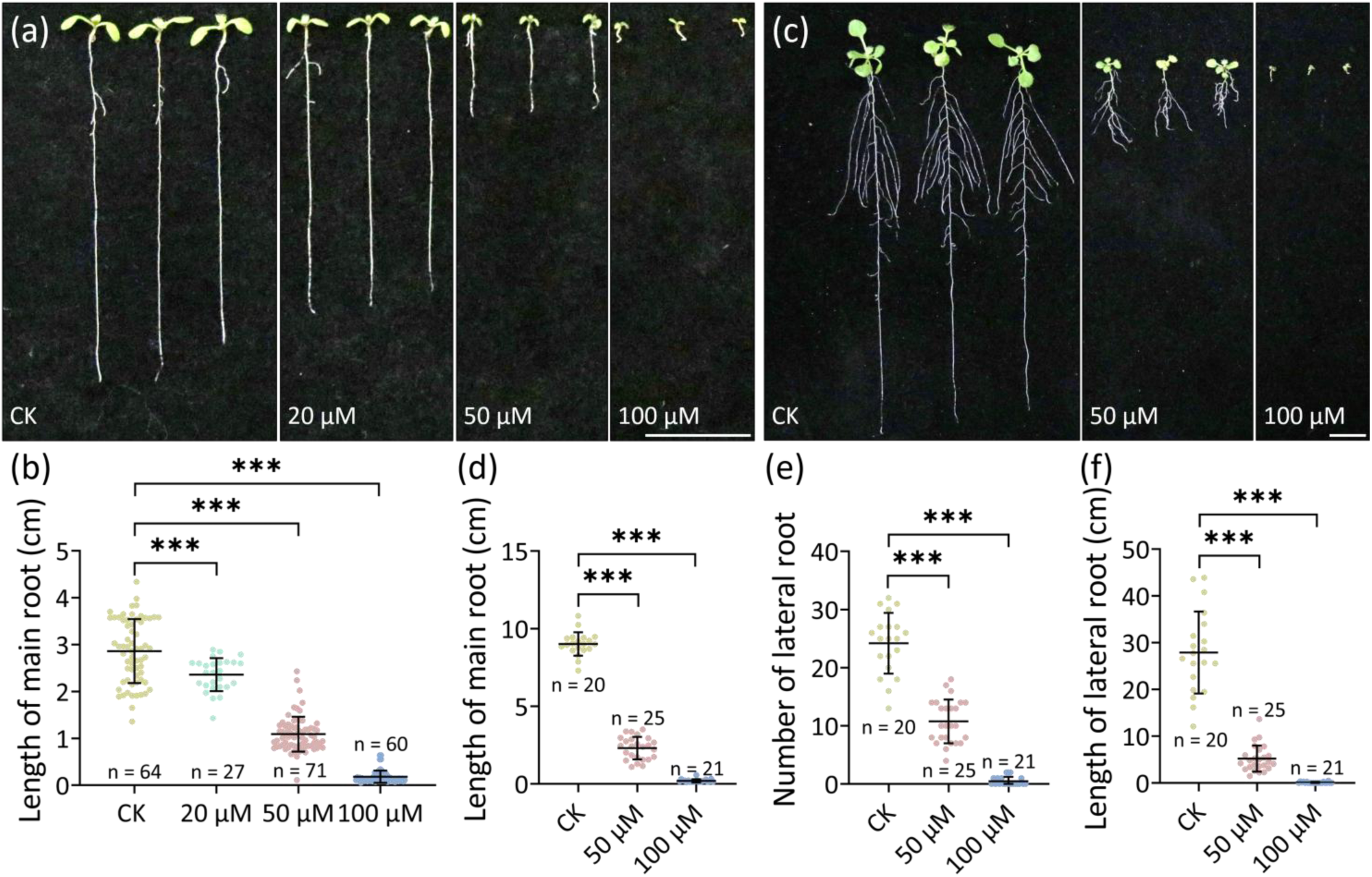
F-53B inhibits root development in Arabidopsis. (a), 8-day-old Col seedlings exposed to F-53B under increasing concentrations. (b), Graph showing the length of main roots in 8-day-old Col seedlings exposed to F-53B. (c), 14-day-old Col seedlings exposed to F-53B under increasing concentrations. (d), Graph showing the length of main roots in 14-day-old Col seedlings exposed to F-53B. (e and f), Graphs showing the number (e) and total length (f) of lateral roots in 14-day-old Col seedlings exposed to F-53B. Significance levels were calculated using unpaired *t* tests; n indicates the number of the analyzed individuals; *** indicates *P* < 0.001. Scale bars, 1 cm.

### 2.2 Preparation of medium with F-53B

The F-53B product, which comprises 97% 6:2 Cl-PFESA and 3% 8:2Cl-PFESA (Wu et al., 2023), was purchased from Jianglai Biotechnology Co., Ltd (Shanghai, China). A 0.25 M F-53B stock solution was prepared and kept at 4°C under dark. The 20, 50 and 100 μM F-53B working solutions, which were chosen by rereferring to (Li et al., 2023b), were prepared using the stock solution in half-strength Murashige & Skoog (1/2 MS) medium. For control medium, 200 μL DMSO was added and mixed in 500 mL 1/2 MS medium.

### 2.3 Preparation of medium with DSB inducers

Camptothecin (CPT) was purchased from Beijing solarbio science & technology co.,ltd (7689-03-4). A 100 mM stock solution A of CPT was prepared, which was then diluted by DMSO into 1 mM as the stock solution B. Finally, a 50 nM working solution of CPT in 1/2 MS medium was prepared. For control medium, 25 μL DMSO was added and mixed in 500 mL 1/2 MS medium. Zeocin was purchased from Life Sciences Solutions Group, Thermo Fisher Scientific (92008). To prepare a 50 ug/mL working solution of zeocin, 250 μL stock solution (100mg/mL) of zeocin was added and mixed in 500 mL 1/2 MS medium.

### 2.4. Generation of the *pMRE11::MRE11-eGFP* and *pMRE11::GUS-eGFP* reporters

To generate the *pMRE11::MRE11-eGFP* reporter, a 2.1 kb coding region fragment of *MRE11* without the stop codon was cloned and fused to the amino-terminal of eGFP via a 15 amino acid linker (GGGGSGGGGTGGGGS), which was then cloned into the binary vector pCB302 (Xiang et al., 1999) under control of the *MRE11* native promoter. To generate the *pMRE11-GUS-eGFP* reporter, the coding sequence of *GUS* was fused to the amino-terminal of eGFP via a 15 amino acid linker (GGGGSGGGGTGGGGS), then cloned into the binary vector pCB302 (Xiang et al., 1999) under control of the *MRE11* native promoter. The construct was transformed into heterozygous *mre11* plants by Agrobacterium tumefaciens (strain GV3101) via the floral dip method (Clough and Bent, 1998). Primers used for plasmid construction are listed in Supplementary material Table S1.

### 2.5. Histochemical staining

To analyze the impact of DSB inducers and F-53B on the activity of the *MRE11* promoter, Arabidopsis seedlings expressing the *pMRE11::GUS-GFP* reporter at the 5^th^ day post vernalization were stained using a GUS staining kit (RTU4032; Real-Times) following the manufacturer’s protocol. After a reaction in the staining solution in dark and 37°C conditions for 24 h, samples were decolorized for 10 days in a ClearSee solution (xylitol [10% (w/v)], sodium deoxycholate [15% (w/v)] and urea [25% (w/v)]) (Kurihara et al., 2015). To evaluate cell viability, 5-day-old Arabidopsis seedlings in 1/2 MS medium with and without F-53B or DSB inducers were stained with a propidium iodide solution (10 μg/mL) for 5 min under dark, and then were washed three times with diluted water. For aniline blue, trypan blue and 4’,6-diamidino-2-phenylindole dihydrochloride (DAPI) staining, 5-day-old Arabidopsis seedlings grown in 1/2 MS medium with and without F-53B were transferred to a aniline blue stain solution (0.1% [m/v] in 0.033% K_3_PO_4_ [m/v]) (Merck, 28983-56-4), trypan blue stain solution (0.4%) (Solarbio, C0040) or a DAPI stain solution (5 μg/mL) (Merck, 28718-90-3), respectively. Seedlings were stained under dark for modulable durations (staining durations for control and F-53B or DSB inducers were kept the same in each single assay), and then were gently washed three times with phosphate buffered solution (1X, 0.01 M, pH = 7.4). For the quantification of aniline blue-stained cell walls, multiple Z-stacks of the seedlings were pictured while the number of cell walls was counted manually.

### 2.6. Live-imaging of reporters

The seedlings of reporters were gently placed on a glass slide immersed in a suitable amount of distilled water, and were then covered by a cover slide. For the *pRPS5A::TagRFP-TUA5* and *pMAP65-3::GFP-MAP65-3* reporters, multiple Z-stacks of seedlings were recorded and the cell walls were counted manually. For the *pCYCB3;1::CYCB3;1-GFP*, *pCDKA;1::CDKA;1-YFP*, *pPIN3::PIN3-GFP*, *pRAD51::RAD51-GFP*, *pUBQ::mCherry-ATG8e*, *p35S::ATG8a-GFP* and *pMRE11::MRE11-GFP* reporters, the seedlings were firstly pictured under a bright-field channel, which was then transferred to a GFP or RFP channel and the seedlings were pictured without changing the Z-stack. In all assays, seedlings under control and chemical-exposure conditions were pictured using the same excitation intensity and the same exposure duration.

### 2.7. Calculation of fluorescence intensity

To measure the fluorescence intensity of CYCB3;1-GFP at the MZ in a seedling, the average fluorescence intensity of the area expressing CYCB3;1-GFP was firstly measured. The average fluorescence intensity of the background of the same picture, i.e., an area besides the seedling, was then calculated and subtracted by the average fluorescence intensity of the CYCB3;1-GFP-expressing area. This method was also used for the measurement of the fluorescence intensity of PIN3-GFP in the columella cells in a seedling. To calculate the fluorescence intensity of CDKA;1-YFP in a seedling, an arrow was drawn from the root cap upwards at the 600 μm position. The fluorescence intensity plot of the section was measured, the values of which subtracted the fluorescence intensity plot of a section besides the seedling. This method was also used for the calculation of the fluorescence intensity of PIN3-GFP in the pericycle zone in a seedling, except for a difference which was that the arrow was drawn from the top of columella cells.

### 2.8. Microscopy

Bright-field and fluorescence images (excitation wavelength peaks from 496 to 553 nm; emission wavelength peaks from 519 to 568 nm) were taken using an Olympus IX83 inverted fluorescence microscope equipped with an X-Cite lamp and a Prime BSI camera. Calculation of fluorescence intensity and bifluorescent images and Z-stacks were processed by Image J.

### 2.9. Statistics

Significance levels were calculated via unpaired *t*-tests or chi-squared (χ^2^) tests using GraphPad Prism (version 8), with the significant level being set as *P* < 0.05. The numbers of analyzed cells or biological replicates are shown in figures or legends; variation bars indicate standard deviation (SD).

## 3. Results

### 3.1. F-53B inhibits root development in Arabidopsis

To determine the impacts of F-53B on plant development, we evaluated seedlings development in the model plant species Arabidopsis (*Arabidopsis thaliana*) by measuring the length of roots of wild-type Columbia-0 (Col-0; ‘Col’ will be used for simplification) exposed to F-53B with elevated concentrations, which have been confirmed with a negative effect on both seed germination and root elongation in wheat (*Triticum aestivum* L.) (Li et al., 2023b; Lin et al., 2020). At the 8^th^ day post vernalization, the length of main roots in Col exposed to increased concentrations of F-53B, i.e., 20, 50 and 100 μM, showed 17.5%, 64.6% and 95.2% reduction, respectively, compared with that in control (Fig. 1a and b). Similar inhibitory effect was observed in Col exposed to 50 or 100 μM F-53B for 14 days (Fig. 1c and d). Besides, F-53B largely reduced the number and total length of lateral roots in Col (Fig. 1c, e and f). These findings reveal a generality of a dosage-dependent impact of F-53B on the development in different plant species.

### 3.2. Root architecture and cell division at the meristematic region are impaired in Arabidopsis exposed to F-53B

To address the mechanism underpinning inhibited root development by F-53B, we stained Col seedlings with aniline blue and measured the length of the meristematic zone (MZ) where cell division occurs, based on the size of cortex cells (Fig. 2a) (Baulies et al., 2022). In control, the length of the MZ in 5-day-old Col seedlings showed an average of 295.2 μm, which was significantly longer than that in seedlings exposed to 50 μM F-53B (Fig. 2b; c, 166.5 μm; e, *P* < 0.001). Under 100 μM F-53B conditions, Col seedlings did not show an obvious MZ (Fig. 2d), which thus were not able to be measured. Meanwhile, Col seedlings exposed to 50 μM F-53B showed a largely reduced number of cortex cells at the MZ than control (Fig. 2f). Moreover, the MZ in Col seedlings under the 50 μM F-53B conditions displayed a significantly smaller number of newly-constructed callosic cell walls marked by bright aniline blue fluorescence (Fig. 2g, h and j, *P* < 0.01), and no newly-formed callosic cell wall was detected in Col exposed to 100 μM F-53B (Fig. 2i). These findings indicated that F-53B suppresses cell division at the MZ, which may contribute to the inhibited root elongation.

**Fig. 2.**
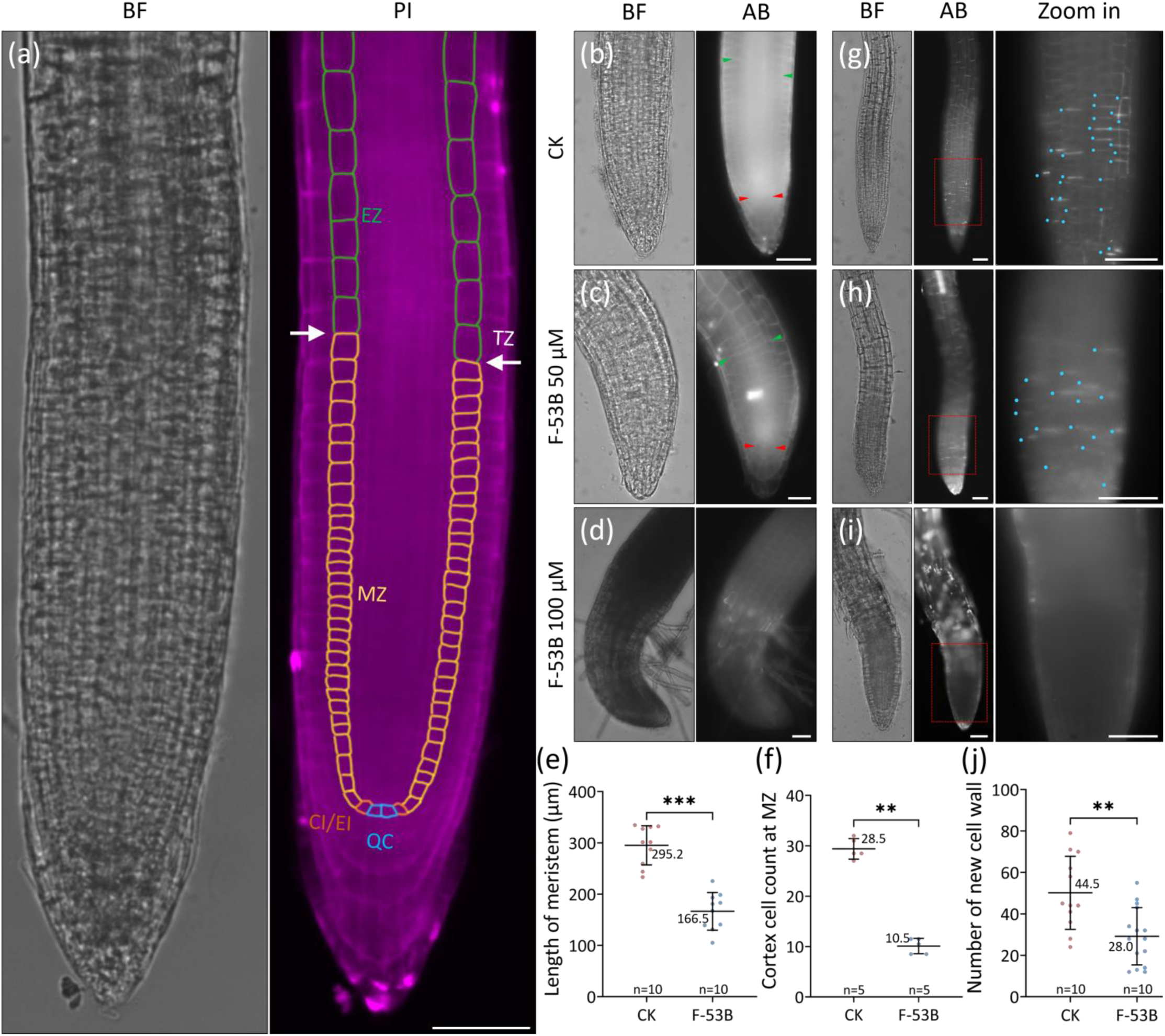
F-53B inhibits meristem development in Arabidopsis seedlings. (a), A PI-stained Col seedling showing different root regions. BF, bright-field; PI, PI-stained; EZ, elongation zone; TZ, transition zone; MZ, meristematic zone; CI/EI, cortex/endodermis initials; QC, quiescent center. The white arrows indicate the border between the MZ and EZ. (b-d), Aniline blue-stained Col seedlings in control medium (b) and exposed to 50 (c) and 100 μM F-53B (d). The green and red arrows point to the top and bottom sites, respectively, of the MZ. AB, aniline blue. (e and f), Graphs showing the length of the MZ (e) and the number of cortex cells at the MZ (f) in Col seedlings under control and F-53B conditions. (g-i), Aniline blue-stained Col seedlings in control medium (g) and exposed to 50 (h) and 100 μM F-53B (i). The red boxes indicate the regions that are zoomed; blue dots indicate newly-formed callosic cell walls. (j), Graph showing the number of newly-formed callosic cell walls in Col seedlings under control and F-53B conditions. Significance levels were calculated using unpaired *t* tests; *** indicates *P* < 0.001; ** indicates *P* < 0.01; n indicates the number of analyzed bio-replicates; values indicate the length of the MZ (e), the numbers of cortex cells (f) or cell walls (j). Scale bars, 50 μm.

### 3.3. F-53B inhibits microtubule-mediated cell plate formation in meristem cells

Microtubule-mediated cell plate formation at anaphase is a key process during cell division (Komaki and Schnittger, 2017). To provide evidence supporting that F-53B prohibits cell division at root tips, we analyzed microtubule organization at the MZ in Col seedlings expressing a *pRPS5A::TagRFP-TUA5* reporter that encodes TUA5 proteins comprising microtubules, and expressing a *pMAP65-3::GFP-MAP65-3* reporter, which encodes MAP65-3 proteins that associate with microtubules and localize at cell plates (Ho et al., 2012; Sofroni et al., 2020). In control, an average of 46.0 cell plates labeled by RFP-tagged TUA5 protein was observed at the MZ (Fig. 3a, b, d and p). In comparison, the seedlings exposed to 50 (Fig. 3f, g, i and p) or 100 μM (Fig. 3k, I, n and p) F-53B showed an average of 24.0 and 8.5 RFP-TUA5-marked cell plates per seedling, respectively. These data indicated that F-53B reduces the microtubule-mediated cell plate formation at root tips. In line with this finding, the number of cell plates labeled by GFP-tagged MAP65-3 proteins was also significantly reduced under 50 (Fig. 3h, j and p; *P* < 0.01) and 100 μM (Fig. 3m, o and p; *P* < 0.001) F-53B conditions. Moreover, seedlings exposed to either concentration of F-53B showed an increased fraction of cell plates with altered orientation or irregular configuration (Fig. 3i, j, m, o, q and r). Overall, these data revealed that F-53B disrupts cell plate formation and microtubule organization at root tips.

**Fig. 3.**
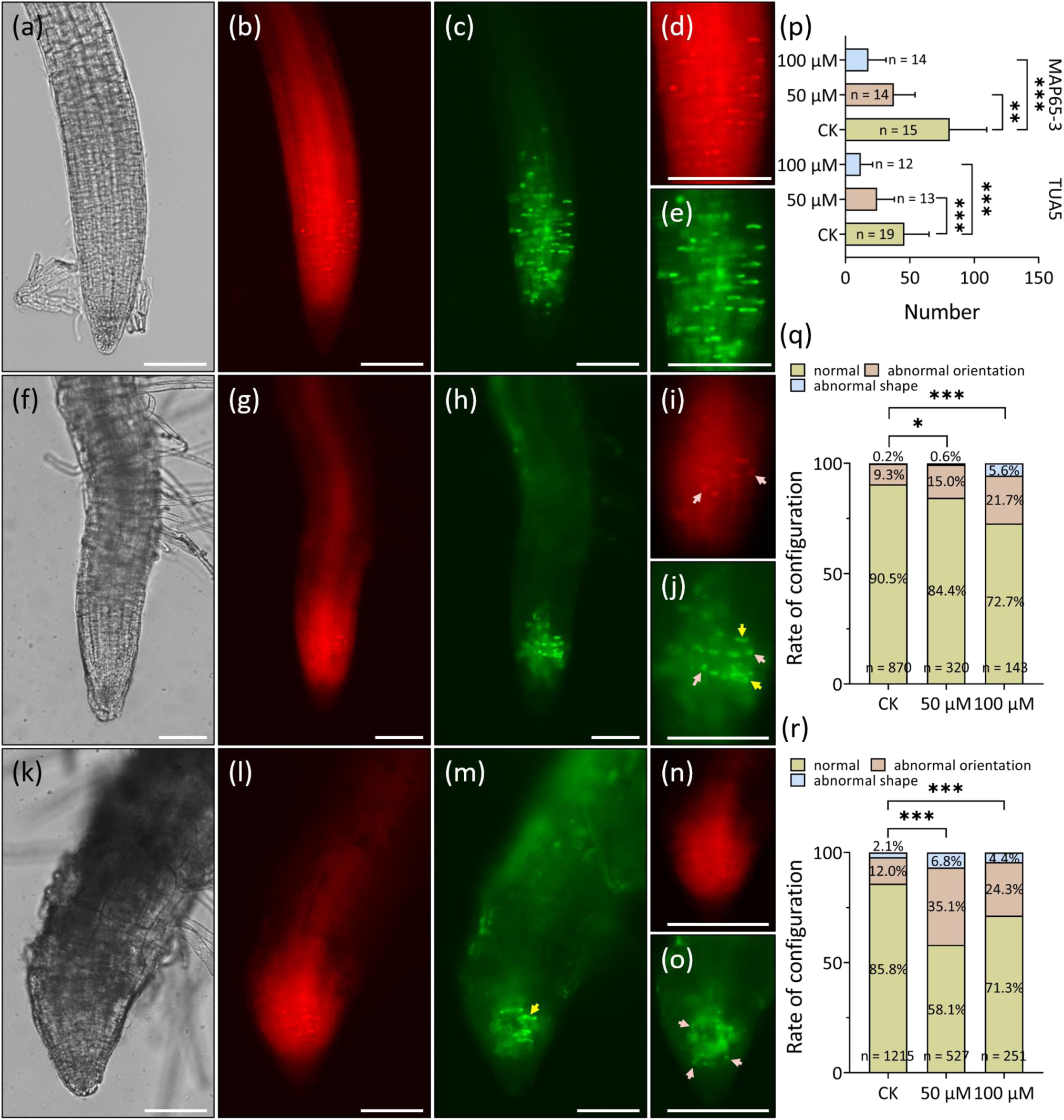
F-53B disrupts cell plate formation and microtubule organization at root tips. (a-o), Col seedlings expressing *pRPS5A::TagRFP-TUA5* (b, d, g, i, l and n) and *pMAP65-3::GFP-MAP65-3* (c, e, h, j, m and o) reporters under bright-field (a, f and k), RFP (b, d, g, i, l and n) and GFP (c, e, h, j, m and o) channels under control (a-e), 50 (f-j) and 100 μM (k-o) F-53B conditions. The white arrows indicate cell plates with altered orientation; yellow arrows indicate cell plates with irregular configurations. (d and e), (i and j) and (n and o) are zoomed-in figures of (b and c), (g and h) and (l and m), respectively. (p), Graph showing the numbers of TagRFP-TUA5 or GFP-MAP65-3-labeled cell plates under control, 50 and 100 μM F-53B conditions. (q and r), Graphs showing the rates of TagRFP-TUA5 (q) or GFP-MAP65-3-labeled (r) cell plates exhibiting different configurations under control, 50 and 100 μM F-53B conditions. Significance levels in (p-r) were calculated based on unpaired *t* tests (p) and χ^2^ tests (q and r); *** indicates *P* < 0.001; ** indicates *P* < 0.01; * indicates *P* < 0.05; frequencies indicate the rates of the corresponding phenotypes; n indicates the number of analyzed bio-replicates (p) or cell plates (q and r). Scale bars, 100 μm.

### 3.4. F-53B reduces the size of the meristematic zone expressing CYCB3;1 and CDKA;1 in Arabidopsis seedlings

Cyclins play a key role in regulating the progression and completion of both mitotic and meiotic cell division (Garrido et al., 2020; Sofroni et al., 2020). Hence, we asked whether F-53B-inhibited cell division is owing to an interfered expression of cyclins. To this end, we analyzed the expression of GFP-tagged B-type cyclin CYCB3;1, which activates the anaphase-promoting complex and regulates spindle organization in both meiosis and mitosis (Garrido et al., 2020; Romeiro Motta et al., 2024), using a *pCYCB3;1::CYCB3;1-GFP* reporter (Sofroni et al., 2020) exposed to 50 or 100 μM F-53B for 5 days. We observed that the recombinant CYCB3;1-GFP protein was mainly expressed at the MZ at root tips (Fig. 4a). However, the size of the area showing CYCB3;1-GFP expression in seedlings of the reporter exposed to 50 or 100 μM F-53B was significantly smaller than that in control (Fig. 4a and b; *P* < 0.05), despite there was no difference in the average fluorescence intensity within the areas between the seedlings under control and F-53B conditions (Fig. 4c).

**Fig. 4.**
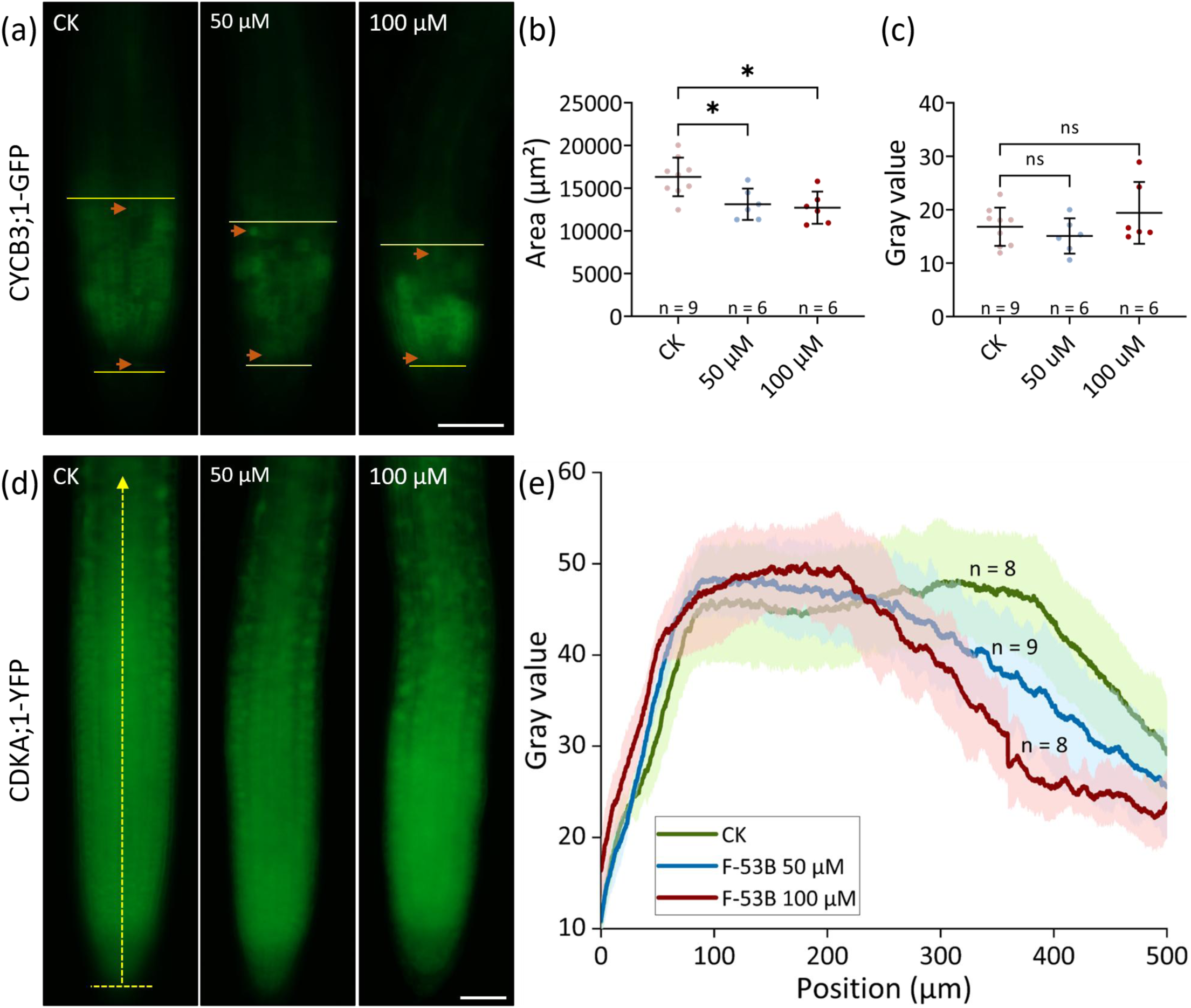
F-53B reduces the size of the meristematic zone expressing CYCB3;1 and CDKA;1. (a), Col seedlings expressing the reporter *pCYCB3;1::CYCB3;1-GFP* under control and F-53B conditions. The yellow lines indicate the regions where the CYCB3;1-GFP fluorescence intensity was calculated; brown arrows indicate the expression of CYCB3;1-GFP. (b), Graph showing the size of area expressing CYCB3;1-GFP in reporter seedlings under control and F-53B conditions. (c), Graph showing the fluorescence intensity of the regions expressing CYCB3;1-GFP in reporter seedlings under control and F-53B conditions. (d), Col seedlings expressing the reporter *pCDKA;1::CDKA;1-YFP* under control and F-53B conditions. (e), Graph showing the fluorescence intensity plot profile of a section indicated by the yellow arrow in seedlings expressing *pCDKA;1::CDKA;1-YFP* under control and F-53B conditions. Significance levels in (b and c) were calculated using unpaired *t* tests; * indicates *P* < 0.05; ns indicates *P* > 0.05; n indicates the number of the analyzed individuals. Scale bars, 50 μm.

Next, we analyzed the impact of F-53B on the expression of the A-type cyclin-dependent kinase, CDKA;1 that plays a crucial role in governing cell cycle in both vegetative and reproductive development (Iwakawa et al., 2006; Nowack et al., 2012). Col carrying the *pCDKA;1::CDKA1-YFP* reporter (Nowack et al., 2006) showed that CDKA;1 was expressed in whole seedlings (Fig. 4d). From the bottom of seedlings, the fluorescence intensity of the CDKA1-YFP recombinant protein increased rapidly and reached peak at the MZ then decreasing upwards (Fig. 4d and e). In seedlings exposed to 50 or 100 μM F-53B, CDKA1-YFP showed a similar expression pattern as control, and no difference in the fluorescence intensity from the bottom to around 200 μm position upwards was detected between control and F-53B conditions (Fig. 4d and e). However, the decrease of the CDKA1-YFP fluorescence intensity occurred earlier in the seedlings exposed to either 50 or 100 100 μM F-53B than that in control, at a position roughly the middle of the region showing the highest fluorescence intensity in control seedlings (Fig. 4d and e). Taken together, these data suggested that F-53B likely does not directly interfere with the expression of CYCB3;1 and CDKA;1, but reduces the length of roots, which subsequently leads to the observed alterations in the size of areas expressing CYCB3;1 or CDKA;1 in seedlings.

### 3.5. F-53B impairs cell viability predominantly at the meristematic zone at root tips

It has been reorted that F-53B induces cell unviability in mammals (Lv et al., 2025; Zhang et al., 2025). To determine if F-53B-impaired root development and cell division are caused by damaged cell viability, we stained Col seedlings exposed to F-53B with trypan blue, which binds to intracellular proteins and marks dead cells. Seedlings in control medium showed a light blue color that specifically showed up on the cell walls (Fig. 5a). In comparison, seedlings exposed to either 50 or 100 μM F-53B exhibited swollen root tips with strong blue color (Fig. 5b-e), indicating that viability of the cells was damaged by F-53B. To further characterize the impacts of F-53B on cell viability, we stained Col seedlings with propidium iodide (PI) that specifically imbeds into DNA in dead cells due to impaired membrane integrity. A positive control was set up by exposing Col seedlings to zeocin, which induces DNA double-strand breaks (DSBs) and thus triggers cell death. Col grown in control medium did not show obvious dead cells at root tips (Fig. 5f), however, 36.7% Col seedlings exposed to zeocin showed PI staining signals above the quiescent center (QC) as shown by previous reports (Fig. 5g and k) (Fan et al., 2022; Yoshiyama et al., 2013). F-53B induced stronger yet, different patterns of PI fluorescence at root tips (Fig. 5g-j). Specifically, under 50 μM F-53B conditions, only a small fraction of seedlings (4.8%) showed a similar PI fluorescence localization as those exposed to zeocin (Fig. 5g and l); while most (81.0%) seedings displayed bright PI fluorescence almost at the entire root tips (Fig. 5h and l). Under 100 μM F-53B conditions, most seedlings (73.4%) showed reduced sizes of area with strong PI fluorescence, which primarily located at the MZ (Fig. 5i, j and l).

**Fig. 5.**
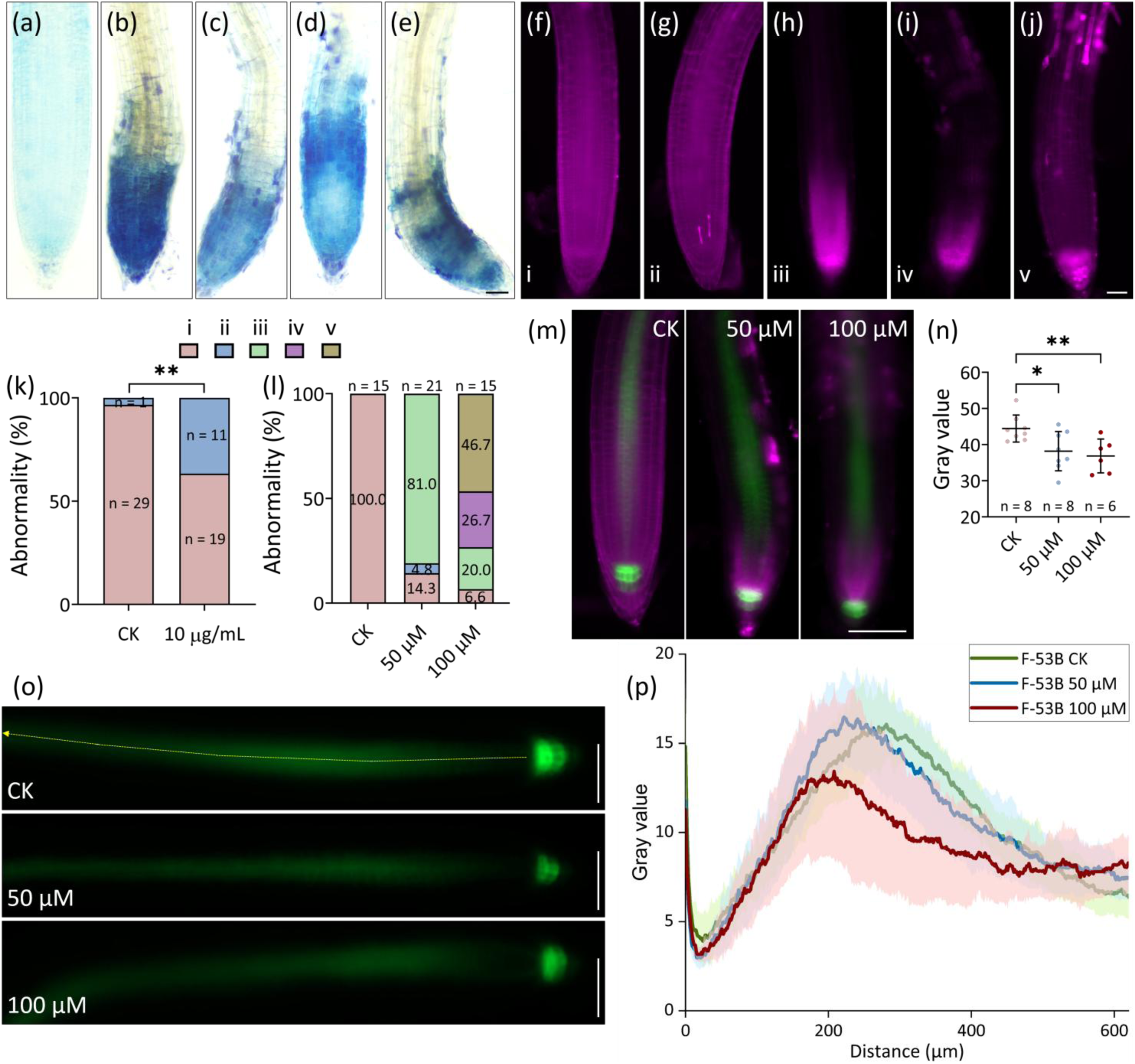
F-53B impairs cell viability primarily at the meristematic zone in seedlings. (a-e), Trypan blue-staining of Col seedlings under control (a), 50 (b and c) and 100 μm F-53B (d and e) conditions. (f-j), Representative images of PI-stained Col seedlings showing different distribution patterns of dead cells under control, or exposed to zeocin or F-53B conditions. (k and l), Graphs showing the fractions of seedlings exhibiting different phenotypes as shown in (f-j) in response to zeocin (k) and F-53B (l). (m), PI-stained Col seedlings expressing *pPIN3::PIN3-GFP* under control, 50 and 100 μm F-53B conditions. (n), Graph showing the PIN3-GFP fluorescence intensity at the columella zone in Col seedlings under control or exposed to F-53B conditions. (o), Col seedlings expressing *pPIN3::PIN3-GFP* under control or exposed to 50 and 100 μm F-53B conditions. (p), Graph showing the fluorescence intensity plot profile of a section indicated by the yellow arrow in Col seedlings expressing *pPIN3::PIN3-GFP* under control, 50 and 100 μm F-53B conditions. Significance levels in (k) and (n) were calculated using χ^2^ tests and unpaired *t* tests, respectively; frequencies in (l) indicate the rates of seedlings showing the corresponding phenotypes; n indicates the number of analyzed individuals; ** indicates *P* < 0.01. Scale bars, 100 μm.

To confirm that F-53B induces cell unviability primarily at the MZ, we analyzed the expression of PIN3, an auxin transpoter protein that is expressed and accumulates in the columella cells below the MZ (Yuan et al., 2021), by performing live-imaging using Col seedlings that express a *pPIN3::PIN3-GFP* reporter (Wang et al., 2021). In control, a strong expression of PIN3 was observed in the columella cells, and it decreased rapidly at the bottom of the pericycle zone followed by an increase and peaking at the EZ, which then decreased gradually upwards (Fig. 5m, o and p). F-53B did not alter the expression and localization patterns of PIN3 in Col seedlings (Fig. 5m, o and p). By refering to the localization of PIN3-GFP at the columella zone, we found that PI fluorescence primarily accumulated at the columella zone and MZ (Fig. 5m). Notably, the PIN3-GFP fluorescence intensity at the columella zone in the seedlings exposed to 50 (Fig. 5m; n, *P* < 0.05; o) or 100 μM F-53B (Fig. 5m; n, *P* < 0.01; o) were significantly lower than that in control. Additionally, the PIN3-GFP fluorescence intensity in the pericycle zone from the EZ upwards in seedlings exposed to 100 μM F-53B was lower than those in control and seedlings exposed to 50 μM F-53B (Fig. 5p). These observations suggested that F-53B impairs cell viability and induces cell death primarily at the MZ, which possibly leads to an interfered PIN3 expression and attenuated auxin transport in seedlings.

### 3.6. F-53B does not increase the expression of RAD51 and MRE11 in root tips

In plants and/or mammals, PFOS and F-53B have been reported to increase the level of phosphorylated histone H2A.X (γH2A.X), which suggest that they induce DNA double-strand breaks (DSBs) (Mah et al., 2010; Pan et al., 2021; Rogakou et al., 1998; Wang et al., 2015). To address whether F-53B induces cell death by triggering ectopic formation of DSBs, we evaluated the impact of F-53B on the expression of RAD51, which plays a key role in DSB repair in both meiotic and mitotic cells. The *pRAD51::RAD51-GFP* reporter, which encodes a recombinant RAD51-GFP protein that assembles at DSB sites thus can be used to indicate DSB amount (Da Ines et al., 2013), was used in our assay. A positive control was set by exposuring the *pRAD51::RAD51-GFP* reporter to the DSB inducer, Camptothecin (CPT) (Li et al., 2023a). As previously reported, the expression of RAD51 was largely increased upon CPT treatment (Fig. 6a and b) (Kutashev et al., 2024). However, the exposure to 50 or 100 μM F-53B did not obviously alter the RAD51-GFP fluorescence intensity at root tips that showed bright PI fluorescence (Fig. 6a, c and d). These findings implied that F-53B does not trigger DSBs in Arabidopsis seedlings.

**Fig. 6.**
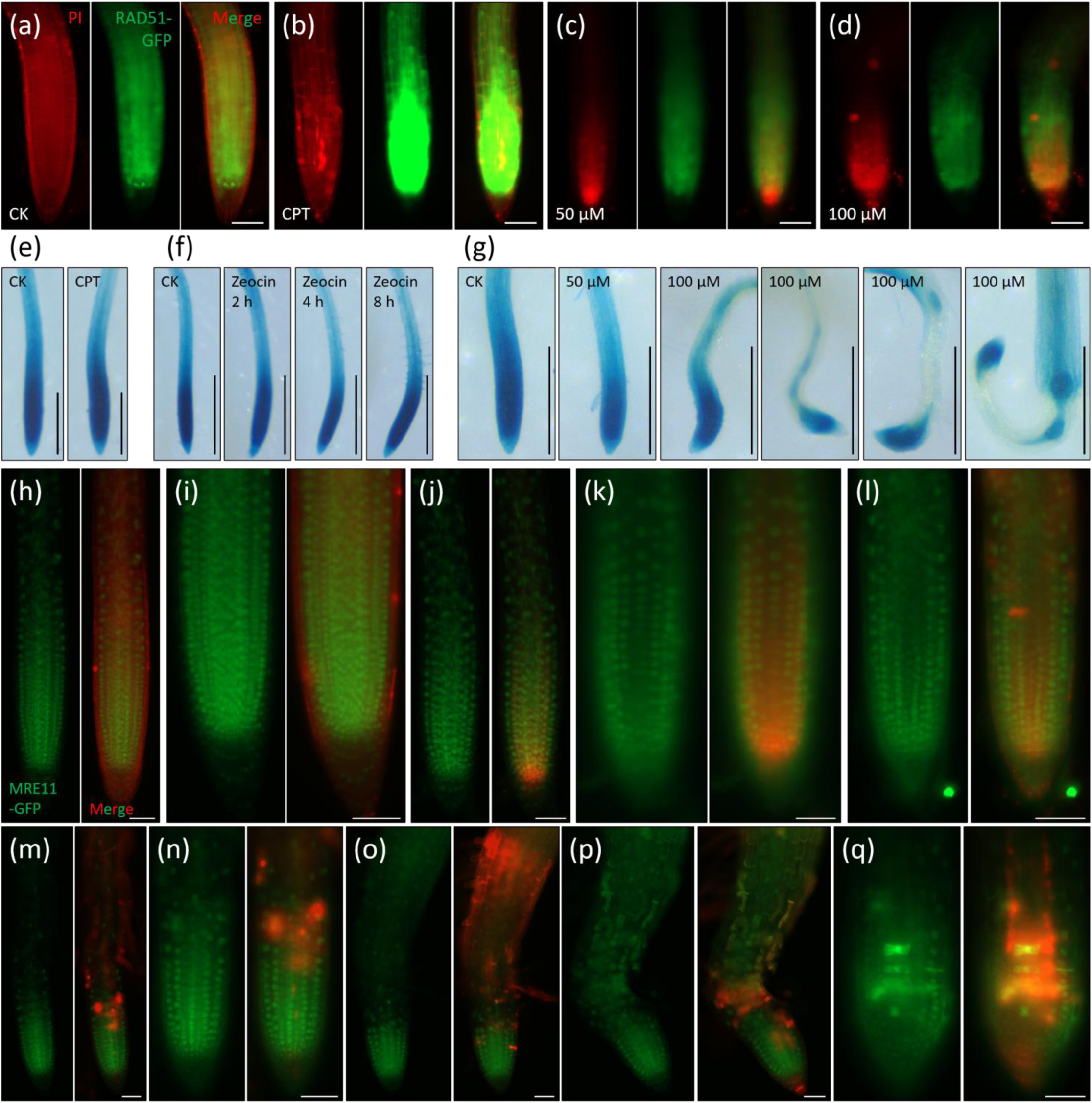
F-53B does not promote RAD51 and MRE11 expression in Arabidopsis seedlings. (a-d), PI-stained *pRAD51::RAD51-GFP* reporter seedlings under control (a), CPT (b), 50 (c) and 100 μM (d) F-53B conditions. (e-g), GUS-staining of *pMRE11-GUS-GFP* reporter seedlings under CPT (e), zeocin (f) and F-53B (g) conditions. (h-q), PI-stained *pMRE11::MRE11-GFP* reporter seedlings under control (h and i), 50 (j-l) and 100 μM (m-q) F-53B conditions. (n) is a closed-up picture of (m). For (a-d and h-q), scale bars, 50 μm; for (e-g), scale bars, 500 μm.

**Fig. 7.**
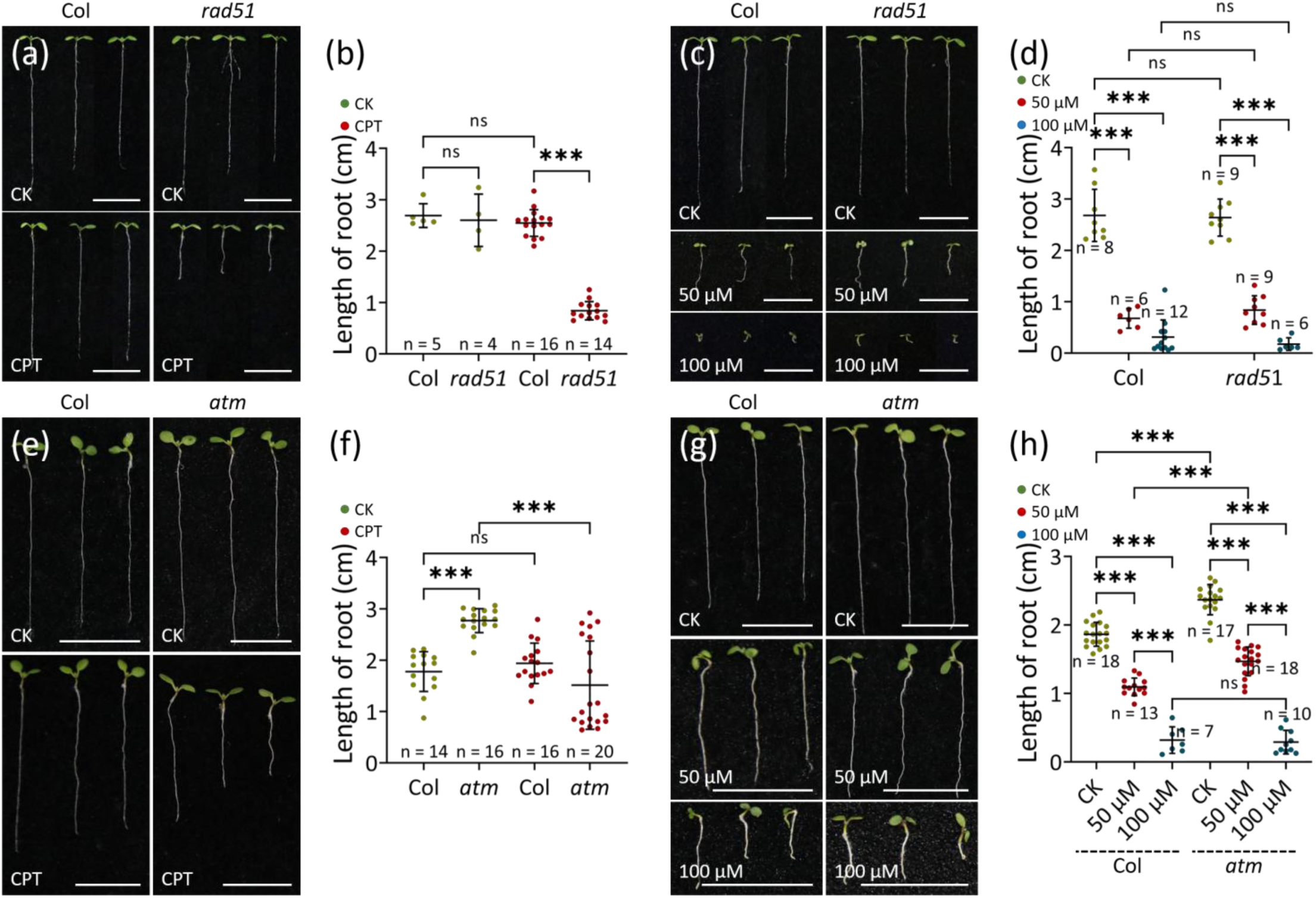
Arabidopsis *rad51* and *atm* mutants show the same response to F-53B as wild-type. (a, c, e and g), 8-day-old seedlings of *rad51* (a and c) and *atm* (e and g) under CPT (a and e) and F-53B (c and g) conditions. (b, d, f and h), Graphs showing the length of seedlings in *rad51* (b and d) and *atm* (f and h) under CPT (b and f) and F-53B (d and h) conditions. Significance levels were calculated based on unpaired *t* tests; n indicates the number of analyzed bio-replicates; *** indicates *P* < 0.001; ns indicates *P* > 0.05. Scale bars, 1 cm.

Next, we analyzed the impact of F-53B on the expression of *Meiotic Recombination 11* (*MRE11*), which plays a conserved role in initiating DSB repair in eukaryotes (Stracker and Petrini, 2011), using a *pMRE11::GUS-GFP* reporter. However, a GUS staining assay revealed that both CPT and another DSB inducer, zeocin, did not obviously alter the blue color at root tips (Fig. 6e and f), hinting that the *MRE11* promoter activity is not responsive to DSBs. Notably, the reporter seedlings exposed to 50 μM F-53B for 5 days showed reduced areas stained with dark blue color, and this effect was pronounced under 100 μM F-53B conditions (Fig. 6g). To further analyze the response of MRE11 to F-53B, we monitored the expression and localization of MRE11 protein in seedlings using a newly-generated *pMRE11::MRE11-GFP* reporter that fully rescued the developmental defect in the *mre11-3* mutant (Supplementary material Fig. S1) (Puizina et al., 2004). In control, dense and circular MRE11-GFP signals were visualized in the whole seedlings, indicating a high expression of MRE11 in the nuclei (Fig. 6h and i). In seedlings exposed to 50 μM F-53B, however, MRE11-GFP signals displayed a slightly sprase localization, especially at the MZ that showed red PI fluorescence (Fig. 6k and l). Reduction and/or impairment of MRE11-GFP expression was more obvious in seedlings exposed to 100 μM F-53B, in which the regular structure of MZ was disrupted (Fig. 6m-q). Since the cells at the region that was not damaged by F-53B and kept normal tissue architecture could express MRE11-GFP regularly, we propose that F-53B-induced defects in MRE11 expression was a consequence of the impaired cell unviability.

### 3.7. Arabidopsis *rad51* and *atm* mutants display the same response to F-53B as the wild-type

To consolidate that F-53B does not induce DSB formation in Arabidopsis seedlings, we searched for genetic evidence by analyzing the root development in Arabidopsis *rad51* mutant in response to F-53B. In a positive control assay, *rad51* exposed to CPT showed a significantly reduced length of seedlings compared with that in control (Fig. 7a and b; *P* < 0.001), which accorded with the notion that RAD51 is essential for root elongation under exogenous DSB inducer treatment (Yu et al., 2023). Under F-53B conditions, the length of seedlings in both Col and *rad51* was largely reduced, which, additionaly, occurrd at similar levels (Fig. 7c and d), indicating that RAD51 is not required for the response of root development to F-53B. We also evaluated the root develompent in Arabidopsis delepted with the conserved kinase Ataxia-Telangiectasia Mutated (ATM), a core regulator of DSB repair through the homologous recombination pathway (Burma et al., 2001; Garcia et al., 2003; Matsuoka et al., 2007). The *atm* mutant behaved the same as *rad51*, which, compared with Col, was more sensitive to CPT but not to F-53B (Fig. 7e-h). These genetic evidence supported that F-53B does not induce DSBs in Arabidopsis seedlings, implying a divergent effect of F-53B on DNA stability across species.

### 3.8. F-53B damages nuclei viability in seedlings

To explore the cause of the impaired cell viability by F-53B, we stained Col seedlings with 4′,6-diamidino-2-phenylindole (DAPI). Under control conditions, Col seedlings showed regularly organized cell walls upon DAPI staining (Fig. 8a), and, at 1-2 min later, DAPI-stained nuclei were visualized in almost all the cells at the MZ and EZ (Fig. 8b and d). Dividing cells at the MZ showed segregating chromosomes at anaphase (Fig. 8c). In seedlings exposed to 50 μM F-53B, cells at root tips showed irregular shape and organization of cell walls (Fig. 8e), and the amount of nuclei stained by DAPI showing fluorescence at the MZ and EZ was reduced (Fig. 8f and g). Notably, differently-sized nuclei in cells at the EZ were observed, which indicated an interfered nuclei stability and/or integrity (Fig. 8g). In seedlings exposed to 100 μM F-53B, cell wall architecture at root tips was severely disrupted, and DAPI-stained nuclei can only be occasionally observed (Fig. 8h and i).

**Fig. 8.**
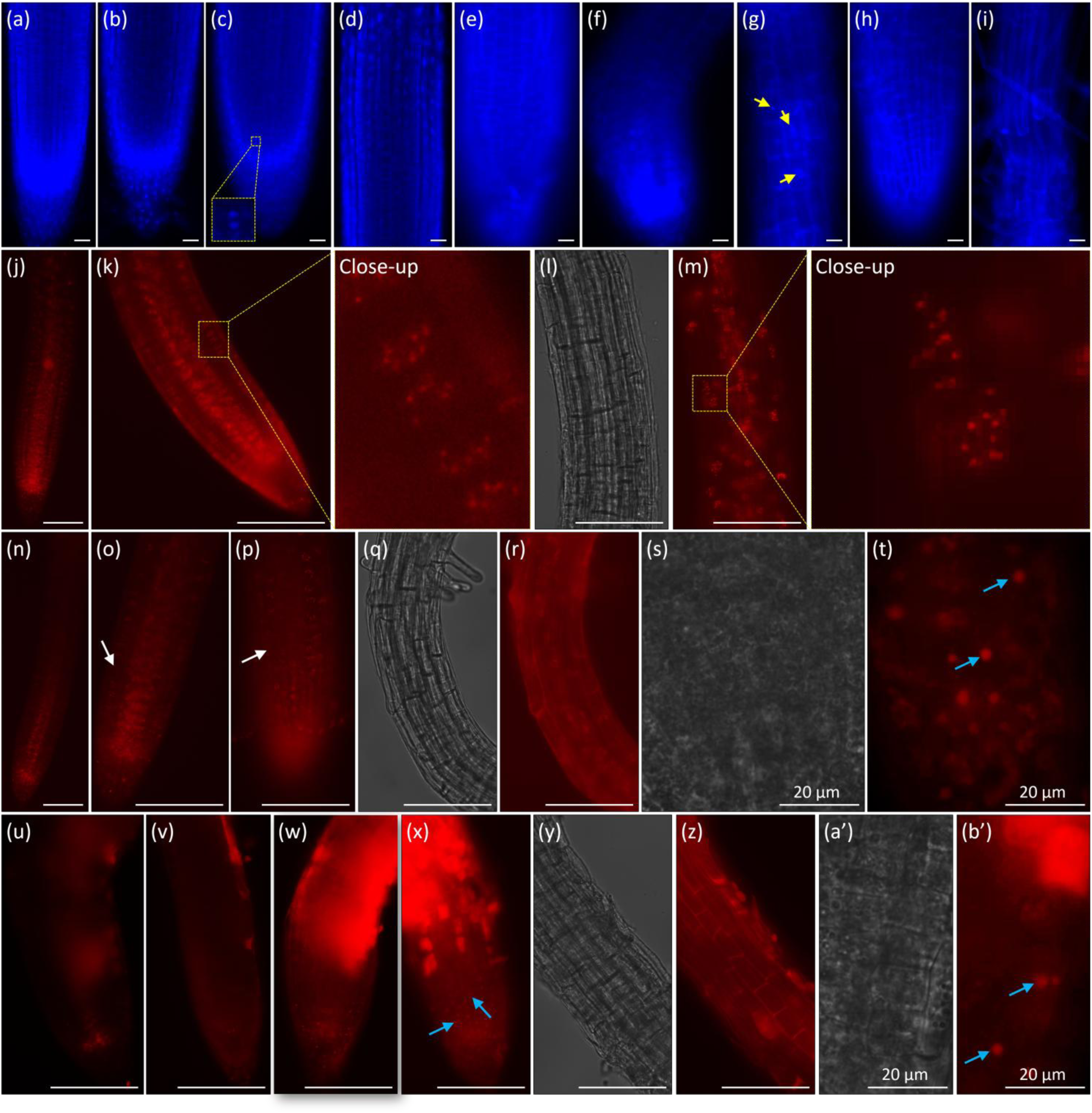
F-53B damages nuclei viability and stability in Arabidopsis seedlings. (a-i), DAPI-staining of Col seedlings under control (a-d), 50 (e-g) and 100 μM (h and i) F-53B conditions. The yellow arrows indicate differently-sized nuclei. (j-b’), Expression of *pCENH3::RFP-CENH3* in Col seedlings under control (j-m), 50 (n-t) and 100 μM (u-b’) F-53B conditions. The white arrows indicate cells without RFP-CENH3 signals; blue arrows indicate irregular RFP-CENH3 signals. For (a-i), scale bars, 20 μm; for (j-r and u-z), scale bars, 100 μm; for (s, t, a’ and b’), scale bars, 20 μm.

To consolidate that F-53B damages nuclei viability and stability, we analyzed the expression of the centromere-specific H3 (CENH3) protein, which lebels and thus allows visualization and quantification of centromeres, using a *pCENH3::RFP-CENH3* reporter (Komaki et al., 2020). Normal expression of the RFP-CENH3 protein can reflect the viability of the nuclei, and the number of its signals can indicate the number of chromosomes. In control seedlings, all the cells at the MZ and EZ yielded RFP-CENH3 signals (Fig. 8j). At the MZ, anaphase-staged cells showing separating RFP-CENH3 signals were visualized (Fig. 8k), and at the EZ, separated nuclei with each nucleus harboring ten RFP-CENH3 signals were recorded (Fig. 8l and m). In comparison, Col seedlings exposed to 50 (Fig. 8n-r, write arrows) or 100 μM F-53B (Fig. 8u-z) displayed less cells expressing RFP-CENH3. Remarkably, irregular and sprase RFP-CENH3 signals, the configuration of which mimicked stress granules that are inducible under abiotic stresses (Li et al., 2024), were observed under both 50 (Fig. 8s and t) and 100 μM F-53B (Fig. 8. x, a’ and b’, blue arrows) conditions. All together, these findings suggested that F-53B damages the viability and stability of nuclei in Arabidopsis seedlings.

### 3.9. F-53B induces ATG8 protein foci formation in seedlings

Cell autophagy maintains intracellular homeostasis and plays a crucial role in mediating stress-induced cellular response and cell death (Liu et al., 2023). To explore whether autophagy is involved in F-53B-induced cell unviability, we analyzed the expression of the ubiquitin-fold protein, Autophagy-Related 8 (ATG8), which is a key regulator of autophagosome formation and has been shown to be responsive to abiotic stress (Zhou et al., 2023). Live-imaging was performed using Col seedlings expressing *pUBQ::mCherry-ATG8e* (Zhuang et al., 2017) or *35S::ATG8a-GFP* (Zhou et al., 2023). In both reporter seedlings exposed to 50 and 100 μM F-53B, cells at the MZ and EZ produced mCherry-ATG8e and ATG8a-GFP puncta, respectively (Fig. 9b-e; h-k), a phenotype not visualized in seedlings under control conditions (Fig. 9a, f and g). These data suggested that F-53B may induce autophagy-mediated cellular responses in Arabidopsis seedlings.

**Fig. 9.**
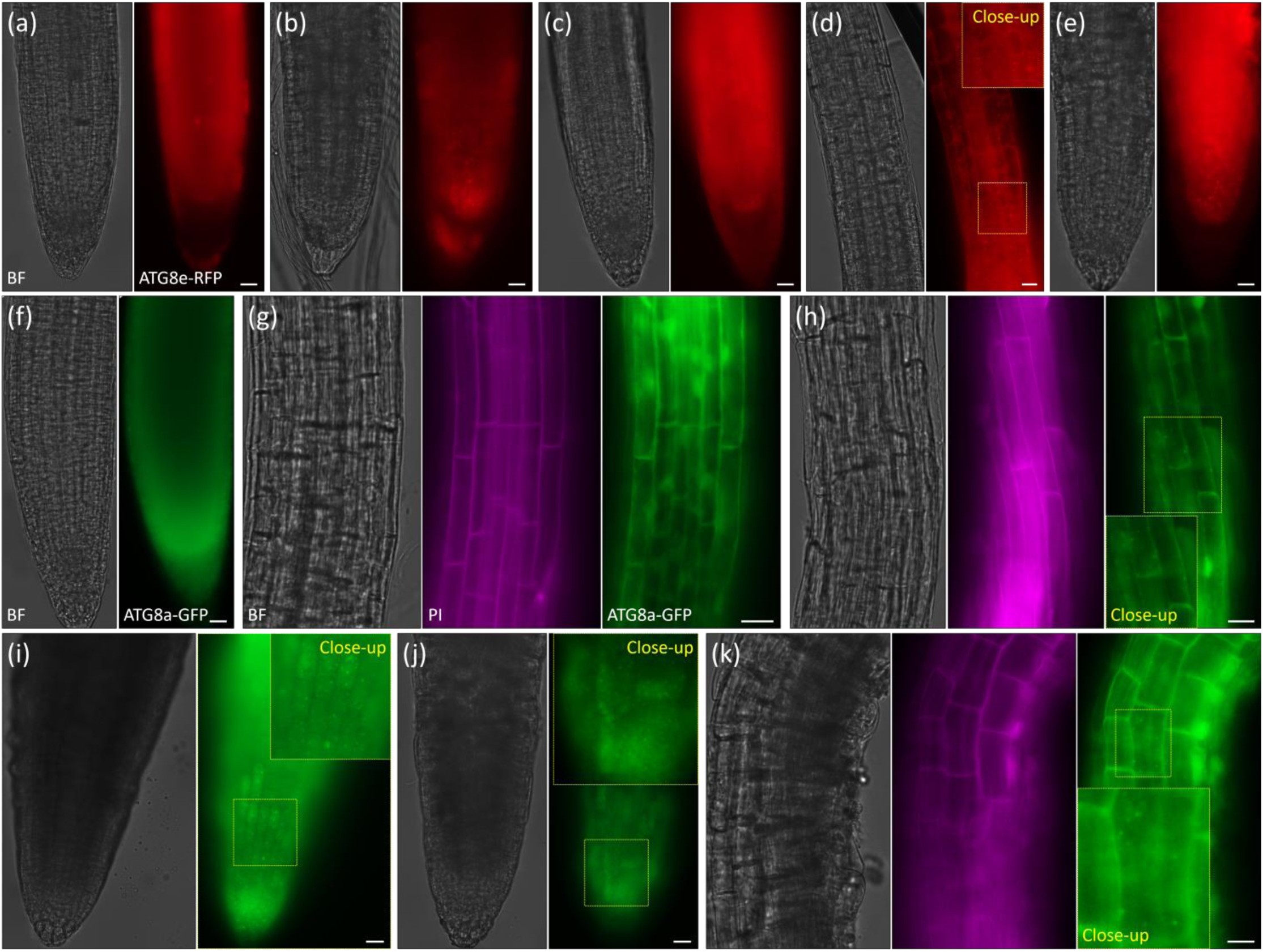
F-53B induces ATG8 protein puncta formation in seedlings. (a-k), Live-imaging of *pUBQ::mCherry-ATG8e* (a-e) and *35S::ATG8a-GFP* (f-k) reporters under control (a, f and g), 50 (b-d, h) and 100 μM (e, i-k) F-53B conditions. Scale bars, 20 μm.

## 4. Discussion

Increasing studies have reported the toxicity of F-53B and PFOA (including PFOS) to development in different plant species (Adu et al., 2023; He et al., 2023; Li et al., 2023b; Pan et al., 2021), however, the mechanisms behind the inhibited plant development by these emerging contaminants were mainly explored through transcriptomic, proteomic and metabolomic studies (Ebinezer et al., 2022; Li et al., 2022; Li et al., 2023b; Lu et al., 2023; Nassazzi et al., 2023; O’Hara and Longstaffe, 2022; Ofoegbu et al., 2022; Pan et al., 2021; Pietrini et al., 2024; Zhang et al., 2022). In the present study, a combination of cytogenetic, molecular and live-imaging microscopy approaches has been applied to dissect the impacts of F-53B on seedling development in *Arabidopsis thaliana*. We report that F-53B prohibits root development and the effect increases along with the elevation of F-53B concentration (Fig. 1), which accords with previous reports (Li et al., 2023b; Lin et al., 2020; Pan et al., 2021; Qu et al., 2010). These facts reveal a dosage-dependent toxicity of F-53B to plants, which is a common feature of the emerging contaminants (Ebinezer et al., 2022; Kim et al., 2024; Pietrini et al., 2024; Rico et al., 2024) that, together with interplays between different environmental factors (Groffen et al., 2023), should be taken into consideration when assessing their impacts on plant development.

We show that F-53B suppresses cell division and induces cell death in Arabidopsis seedlings (Fig. 2; Fig. 3; Fig. 5), which are likely the key cellular defects that result in impaired root elongation in plants. Since F-53B-suppressed cell division and cell death predominantly occur at the MZ, the defects in cell division could be a consequence of the impaired cell viability. The hypersensitivity of the MZ cells to F-53B suggests that F-53B is especially toxic to cells undergoing nuclei division or a high level of metablic activities that need active energy metabolism. In support of this hypothesis, both F-53B and PFOA (including PFOS) have been found to impose strong impacts on biosynthetic processes and the metabolism of amino acids and carbohydrates (Li et al., 2021; Li et al., 2023b; Ofoegbu et al., 2022). We show that the formation and configuration of cell walls at the MZ in Arabidopsis seedlings are prohibited and disorganized by F-53B (Fig. 2), likely resulting from an impairment of the formation and organization of the microtubule-mediated cell plate (Fig. 3). In mammals, PFOS and PFAS disrupt actin cytoskeleton organization in Sertoli cells, at least partially by interfering with the localization of the branched actin polymerization protein Actin-Related Protein 3 and the actin bundling protein Palladin, which consequently leads to truncated actin microfilament and damaged cell morphology and function (Gao et al., 2017; Wan et al., 2020). Moreover, in maize and zebrafish, PFAS (including PFOS) influences the expression of proteins involved in microtubule cytoskeleton organization (Ebinezer et al., 2022; Satbhai et al., 2025).

Based on the findings in our study and these reports, we propose that cytoskeleton network is a main target of the emerging contaminants. It is pssible that F-53B disrupts cytoskeleton organization in Arabidopsis seedlings via the similar mechanism as PFOS does, i.e., by altering the expression, localization and/or function of the cytoskeleton regulators and/or components.

In both mammals and plants, F-53B has been found to increase the level of ɤH2A.X in somatic cells, suggesting that F-53B triggers DSB formation in organisms (Li et al., 2023b; Pan et al., 2021; Qiu et al., 2024). In this study, however, we found that the expression of RAD51 is responsive to DSB inducer but not to F-53B, and, the root development in the *rad51* and *atm* mutants shows the same F-53B sensitivity as that in wild-type (Fig. 6; Fig. 7). These molecular and genetic data suggest that F-53B does not induce DSBs in Arabidopsis seedlings, which thus demonstates that F-53B-induced cell unviability in Arabidopsis seedlings is not caused by ectopic DSB formation, and suggests that F-53B has divergent impacts on DNA stability across species. The Arabidopsis seedlings exposed to F-53B show varied sizes of nuclei; in addition, nuclei in numerous cells were failed to be stained by DAPI and failed to express the CENH3 reporter (Fig. 8). These data indicate that nulcei stability and/or function in those cells are damaged, which could be the main cause of the F-53B-induced cell unviability in seedlings. F-53B induces oxidative stress in wheat roots and shoots, and, in mung bean roots, F-53B triggers production of reactive oxygen species (ROS), hydroxyl radical (·OH) (Li et al., 2023b; Lin et al., 2020; Pan et al., 2021). Hence, we speculate that F-53B-damaged nuclei stability, viability or integrity in Arabidopsis seedlings could at least partially be owing to an ROS-induced programmed cell death and/or apoptosis (Petrov et al., 2015). This study, however, has not answered which kind of DNA damage is triggered by F-53B, which is a particular important question to be addressed in future studies.

Moreover, we found that F-53B alters the expression of the auxin transporter PIN3, suggesting that the content and/or transport of auxin in seedlings is interfered by F-53B (Fig. 5). PFOA and PFOS have been reported to inhibit root growth in Arabidopsis by modulating auxin signalling pathways, and ABA plays a role in the inhibition (Zhang et al., 2022). Hence, we propose that F-53B prohibits root development in Arabidopsis at least partially via the same mechanism as PFOA and PFOS do, which is by perturbing the metabolism and/or signalings of phytohormones. Furthermore, we show that F-53B induces ATG8 protein foci formation in seedlings, suggesting that F-53B evokes autophagy-mediated cellular responses, which may subsequently trigger programmed cell death that facilitates cellular homeostasis, and/or activate stress responses (Zhou et al., 2023). Taken together, our study provides cytogenetic and molecular insights into the tocixity of F-53B to plant cells, which paves a road for further characterizing the mechanisms mediating cellular response and adaption to F-53B in plants.

## 5. Conclusion

In this study, we report that F-53B inhibits root development in a dosage-dependent manner in Arabidopsis. F-53B impairs microtubule-mediated cell plate formation, cell wall organization and cell division in seedlings. Histochemical staining reveals that F-53B damages cell viability predominantly at the meristematic zone. Gene expression and genetic studies demonstrate that F-53B does not induce DNA double-strand breaks (DSBs) in Arabidopsis seedlings, suggesting a divergent impact of F-53B toxicity on DNA stability across species. DAPI staining and live-imaging analysis of CENH3 expression reveal that F-53B disrups nuclei viability and stability at root tips, which likely results in cell unviability. Moreover, we show that F-53B induces ATG8 protein foci formation at both the meristematic and elongation zones, suggesting that F-53B triggers autophagy-mediated cellular responses. Overall, this study provides cytological and molecular insights into the toxicity of F-53B to plant cells. Future studies should address: 1), which kind of DNA damage is triggered in plant cells by F-53B, and how? 2), the genetic and molecular factors and signalings that mediate the perception, uptake and responses to F-53B in plants.

## CRediT authorship contribution statement

J.Z., W.P., Y.C., X.C., H.F., Y.L., Z.C., N.H. and Y.H. performed experiments; Y.Q., G.Y., Z.R. and W.W. contributed to data analysis; X.L. and B.L. conceived project, analyzed data, wrote and edited the manuscript. All authors have read and agreed with the manuscript prior to submission.

## Declaration of Competing Interest

The authors declare that they have no known competing financial interests or personal relationships that could have appeared to influence the work reported in this paper.

## Data Availability

The data that support the findings of this study are available from the corresponding author (B.L.) upon reasonable request.

## Acknowledgements

This study was supported by National Natural Science Foundation of China (22206127 to X.L., 32101571 to Z.R., 32270364 and 31270361 to YG), Hubei Provincial Natural Science Foundation (2024AFB695 to B.L.), the Program of Science and Technology Commission of Shanghai Municipality (21ZR1435800 to X.L.), the Fundamental Research Funds for the Central Universities, South-Central Minzu University (CZZ24011 to B.L., CZQ24012 to W.W., and CZZ21004 and CZZ24011 to G.Y.), Zhejiang Provincial Natural Science Foundation (ZCLTGN24C1601 To Z.R.), the Zhejiang Sci-Tech University Start-up Fund (22052138-Y To Z.R.), and the fund for the Research Center for Biotechnology Application in College of Life Sciences, South-Central Minzu University (XTZ24021). The authors thank Arp Schnittger (UH) for sharing the *pCENH3::RFP-CENH3*, *pCYCB3::CYCB3-GFP*, *pCDKA;1::CDKA1-YFP*, *pMAP65-3::GFP-MAP65-3* and *pRPS5A::RFP-TUA5* reporters; they thank Charles I White and Olivier Da Ines (iGReD) for sharing the *pRAD51::RAD51-eGFP* reporter; they thank Liwen Jiang (CUHK) for sharing the *pUBQ::mCherry-ATG8e* reporter; they thank Caiji Gao (SCNU) for sharing the *p35S::ATG8a-GFP* reporter; they thank Erfang Kang (HNU) and Jirong Huang (SNU) for sharing the *pPIN3::PIN3-GFP* reporter.

## Appendix A. Supplementary material

Supplementary data associated with this article can be found in the online version.

## Notes

### Competing Interest Statement

The authors have declared no competing interest.

